# Different DGAT1s show different TAG synthesis abilities and a specific amino acid substitution enhances oil accumulation

**DOI:** 10.1101/2021.09.19.461007

**Authors:** Tomoko Hatanaka, Yoshiki Tomita, Daisuke Matsuoka, Daisuke Sasayama, Hiroshi Fukayama, Tetsushi Azuma, Mohammad Fazel Soltani Gishini, David Hildebrand

**Affiliations:** Graduate School of Agricultural Science, Kobe University, Kobe, Hyogo, Japan, 675-8501; Department of Production Engineering and Plant Genetics, Faculty of Sciences and Agricultural Engineering, Campus of Agriculture and Natural Resources, Razi University, Kermanshah, Iran; Department of Plant and Soil Sciences, University of Kentucky, Lexington, KY, USA, 40546-0312

**Keywords:** Acyl-CoA:diacylglycerol acyltransferase, *Arabidopsis thaliana*, DGAT1, triacylglycerol, site-directed mutagenesis, *Vernonia galamensis*, yeast strain H1246

## Abstract

Triacylglycerols (TAGs) are the major component of plant storage lipids. Acyl-CoA:diacylglycerol acyltransferase (DGAT) catalyzes the final step of the Kennedy pathway, and responsible for plant oil accumulation. We previously found DGAT activity of *Vernonia galamensis* DGAT1 was distinctively higher than that of *Arabidopsis thaliana* DGAT1 and soybean DGAT1 in a yeast microsome assay. In this study, the *DGAT1* cDNAs of Arabidopsis, Vernonia, soybean, and castor were introduced into Arabidopsis (ecotype Col-0). All Vernonia *DGAT1* expressing lines showed a significantly higher oil content (average 49% relative increase compared to the wild type) followed by soybean, and castor. Most Arabidopsis *DGAT1* over-expressing lines did not show a significant increase. In addition to these four *DGAT1*s, sunflower, Jatropha and sesame *DGAT1* genes were introduced into the TAG biosynthesis defective yeast mutant (H1246). In the yeast expression culture, DGAT1s from Arabidopsis, castor, and soybean only slightly increased TAG content, however, DGAT1s from Vernonia, sunflower, Jatropha, and sesame remarkably increased TAG content more than 10 times higher than the former three DGAT1s. Three amino acid residues were characteristically common in the latter four DGAT1s. Using soybean *DGAT1*, these amino acid substitutions by site-directed mutagenesis was performed and analyzed. These substitutions substantially increased the TAG content.

**Highlight:** *DGAT1s* from several plant species were tested their TAG accumulation promotion in Arabidopsis and yeast. They were divided into high and low function and single amino acid substitution enhanced function

## Introduction

Vegetable oil is one of the most important renewable resources in the world. In addition to a wide range of food applications, vegetable oils are also valuable as renewable chemical derivatives such as lubricants, paints, adhesives, varnishes, plasticizers, and biodiesel (Carlsson, 2009; 2011; Jaworski and Cahoon, 2003; Ohlrogge, 1994). Triacylglycerol (TAG) is a major component of plant storage lipids and is synthesized by continuous esterification of acyl chains from acyl-CoA at the *sn*-1, -2, -3 positions of the glycerol backbone (Ohlrogge and Browse, 1995).

As enzymes that catalyze the final stage of this TAG synthesis, three types of enzymes were identified; acyl-CoA: diacylglycerol acyltransferase (DGAT) (Cases *et al*., 1998; Hobbs *et al*., 1999; Zou *et al*., 1999), phospholipid: diacylglycerol acyltransferase (PDAT) ((Dahlqvist *et al*., 2000) (Oelkers *et al*., 2000; Stahl *et al*., 2004) and diacylglycerol transacylase (DAG transacylase) (Fraser *et al*., 2000; Stobart *et al*., 1997). Of these enzymes, DGAT is the most important and has been suggested to be the rate-determining factor in TAG synthesis. In eukaryotes, several classes of DGATs have been identified based on differences in structure and activity (He *et al*., 2004; Lung and Weselake, 2006) DGAT1 belongs to the membrane-bound O-acyltransferase (MBOAT) family (Cases *et al*., 1998), and is usually predicted to have 9 or 10 transmembrane domains in higher plants. The gene encoding *DGAT1* has been isolated and characterized from many plant species (Banilas *et al*., 2011; Beisson *et al*., 2003; Bouvier-Nave *et al*., 2000; Hatanaka *et al*., 2003; Hobbs *et al*., 1999; Jako *et al*., 2001; Li *et al*., 2013; Li *et al*., 2010a; Nykiforuk *et al*., 2002; Shockey *et al*., 2006; 2008; Yu *et al*., 2006).

The DGAT2 protein has only a few predictive transmembrane domains and belongs to the monoacylglycerol acyltransferase (MGAT) family. Both DGAT1 and DGAT2 are endoplasmic reticulum (ER) membrane-binding enzymes, however, their amino acid sequences are not homologous to each other (Cases *et al*., 2001; He *et al*., 2004; Kroon *et al*., 2006; Lardizabal *et al*., 2001; Shockey *et al*., 2006; Xu *et al*., 2013). DGAT2 has been reported to contribute to the accumulation of unusual fatty acids that contain hydroxyl and epoxy groups (Burgal *et al*., 2008; Kroon *et al*., 2006; Li *et al*., 2010b). DGAT1 and DGAT2 are ER membrane-bound, while the third type, DGAT3, is a soluble cytoplasmic type identified in peanuts (*Arachis hypogaea*) (Saha *et al*., 2006) and *Arabidopsis thaliana* (Hernandez *et al*., 2012). A fourth type of DGAT has a sequence similar to DGAT2 and has both wax ester synthase (WS) and DGAT activity (WS / DGAT) (Kalscheuer and Steinbuchel, 2003; King *et al*., 2007; Li *et al*., 2008; Rosli *et al*., 2018; Xu *et al*., 2021). The most common DGAT is type 1 (DGAT1), which contributes to most of the TAG synthesis (Sanjaya *et al*., 2013). Many studies have reported that increased expression of the *DGAT1* gene increases oil content (Andrianov *et al*., 2010; Lardizabal *et al*., 2008; Rao and Hildebrand, 2009; Taylor *et al*., 2009).

*Vernonia galamensis* is an annual herbaceous plant native to East Africa belonging to the Asteraceae family. It contains about 40% oil in its seeds. *V. galamensis* is known to an epoxy fatty acid accumulator (Ayorinde *et al*., 1988; Carlson and Chang, 1985; Perdue *et al*., 1986). Vernonia seed oil contains up to 80% vernolic acid (*cis*-12,13-epoxy-*cis*-9-octadecenoic acid) (Anderson *et al*., 1993; Neff *et al*., 1993) From Vernonia developing seeds, two types of *DGAT1* genes (*VgDGAT1A* and *VgDGAT1B*) have been isolated and the various characteristics have been investigated (Hatanaka *et al*., 2003; Yu *et al*., 2008). Expression of the *Stokesia laevis* epoxygenase gene in soybean seeds showed undesired phenotypic changes in transformed seeds such as wrinkles and weight reduction, however, these negative side effects were restored by co-expression with *VgDGAT1A* (Li *et al*., 2012). Even though Vernonia DGAT2 is more effective than DGAT1s vernolic acid accumulation (Li *et al*., 2010b), VgDGAT1A also significantly increased vernolic acid accumulation in soybean seeds.

In our previous study, in a microsome assay using an expression system in wild-type yeast (*Saccharomyces cerevisiae*), Vernonia DGAT1 showed about 40 times more active than Arabidpsis DGAT1 and about 9 times more active than soybean (*Glycine max*) DGAT1 (Hatanaka *et al*., 2016). However, in this system using wild-type yeast, the activity of *A. thaliana DGAT1* was close to that of the vector control. Therefore, it is unclear that the activity of DGAT1 expressed in wild-type yeast may or may not match these results when they are expressed in plants.

In this study, the effectiveness of DGAT1s among plant species was investigated. The *DGAT1* genes evaluated were derived from four species, *V. galamensis*, soybean (*Glycine max*), castor (*Ricinus communis*), and *A. thaliana* for overexpression, were introduced into Arabidopsis, and their effects on the seed oil content and fatty acid composition were investigated. Further, another three *DGAT1* cDNAs were obtained from sunflower (*Helianthus annuus*), Jatropha (*Jatropha curcas*), and sesame (*Sesamum indicum*) and these seven DGAT1 genes including those above were expressed in the yeast quadruple mutant H1246 strain (*S. cerevisiae*) (Sandager *et al*., 2002), which is deficient in synthesizing TAG and their TAG contents were measured. The amino acid sequence of sunflower DGAT1 is very close to Vernonia DGAT1s and there are reports that Jatropha *DGAT1* (Misra *et al*., 2013) and sesame *DGAT1* (Wang *et al*., 2014) are effective for increasing seed oil contents of transgenic *A. thaliana* lines expressing these genes. On the other hand, direct protein engineering has been used to generate DGAT1 variants with enhanced activity (Roesler *et al*., 2016; Siloto *et al*., 2009). Chen *et al*. (2017) reported that a single amino acid residue substitution leads to enhance *B. napus* DGAT1 activity in yeast. Here, we aim to clarify the diversity of DGAT1 properties among plant species, and find factor(s) to increase their effectiveness in TAG accumulation.

## Materials and methods

### DGAT1 cDNA cloning and the construction of plant and yeast expression vector

Total RNAs were extracted from developing seeds of *A. thaliana* (ecotype, Col-0), soybean (cv. Jack), castor, Jatropha, and sesame (cv. Kanto-1) using an RNeasy Plant Mini Kit™ according to the manufacturer’s instructions (Qiagen, https://www.qiagen.com). Reverse transcription of RNAs were carried out using PrimeScript II 1^st^ strand cDNA Kit™ (Takara, Japan). The coding sequences of *DGAT1s* were amplified by a high-fidelity KOD-Plus-Neo polymerase (Toyobo, Japan) using end-specific primers containing restriction sites. The GenBank numbers of DGAT1 genes in this study are listed in Supplementary Table S1. The primers used in this section are listed in Supplementary Table S2-1. Seeds of sesame (cv. Kanto-1) were gifted from the NARO (National Agriculture and Food Research Organization) Genebank, Japan..

### Arabidopsis transformation and growth condition

For plant expression the amplification products, *AtDGAT1, GmDGAT1A*, and *RcDGAT1* were subcloned into the respective sites of the pPHI4752 vector containing a phaseolin promoter, which confers strong seed-specific expression of transgenes (Bustos *et al*., 1998; Kawagoe *et al*., 1994). The phaseolin promoter cassette containing the coding region of each target gene was transferred into the corresponding sites of the binary pCAMBIA1301, T-DNA vector (Cosmo Bio, Japan). Regarding *VgDGAT1A*, the vector constructed in the previous study (Li *et al*., 2012) was used.

The recombinant plasmids were transformed into *Agrobacterium tumefaciens* strain EHA105 by the freeze-thaw method. Transformation of *A. thaliana* (ecotype Col-0) plants was carried out using the floral dip method (Clough and Bent, 1998). T1 seeds were grown on a solid selection medium which is composed of MS salts (Murashige and Skoog, 1962), B5 vitamins (Gamborg *et al*., 1968), 1% (w/v) sucrose, 25 mg/L (w/v) hygromycine, and 0.8% agar. T2 seeds were harvested and the seed checked for total lipid and TAG contents. Two individual lines that contained higher seed oil levels were selected and grown at 23°C under long-day conditions (16 h of light/8 h of dark). The developing siliques of T2 generations were used for qPCR and mature T3 seeds were used for lipid analyses.

### Quantitative RT-PCR

The expression of *DGAT1*s in the transgenic Arabidopsis lines was analyzed by quantitative RT–PCR (qPCR) on a Thermal Cycler Dice Read Real-Time System (Takara, Japan) with the SYBR Green Premix EX Taq GC™ (Takara, Japan) according to the manufacture’s instruction with the primers shown in Supplementary Table S2-2. Total RNA was isolated from immature siliques using the ISOSPIN Plant RNA Kit™ (Nippon Gene, Japan), which was used to synthesize first-strand cDNA using the SuperScript II first-strand cDNA synthesis Kit™ (Takara, Japan). The Arabidopsis *Actin2* gene was used as the internal reference gene.

### Yeast transformation and culture

For the yeast expression system, coding regions of *DGAT1*s (*AtDGAT1, RcDGAT1, GmDGAT1A, VgDGAT1A, HanDGAT1, JcDGAT1*, and *SiDGAT1*) were cloned into the yeast vector pYES2 (Invitrogen). *S. cerevisiae* strains were then transformed with the experimental constructs using the lithium acetate mediated method as well as an empty vector control. The *S. cerevisiae* strains used in this study were the quadruple knock-out strain H1246 (*MATa are1-D::HIS3, are2-D:: LEU2, dga1-D::KanMX4 and lro1-D::TRP1 ADE2*), containing knockouts of all four neutral lipid biosynthesis genes *DGA1, LRO1, ARE1* and *ARE2*, kindly distributed by Dr. S. Stymne (Sandager *et al*., 2002). All transformants were selected on synthetic complete medium lacking uracil (SC-U, 6.7% (w/v) of Yeast Nitrogen Base, 0.01% (w/v) of adenine, arginine, cysteine, leucine, lysine, threonine and tryptophane, 0.050% (w/v) of aspartic acid, histidine, isoleucine, methionine, phenylalanine, proline, serine, tyrosine and valine) supplemented with 2% (w/v) glucose. The recombinant yeast cells were first cultivated in liquid minimal medium containing 2% (w/v) galactose for one day, and then inoculated in a 50 mL induction medium [SC-U, 2% (w/v) galactose, and 2% (w/ v) glucose] in 200-mL shaking Erlenmeyer flasks with baffles at an initial optical density of 0.4 at 600 nm (OD_600_) according to Invitrogen’s instructions. The induction culture was performed at 30°C with shaking at 180 rpm for five days. The expression of transgenes was checked by reverse transcription-PCR. Total RNA extraction and the first strand cDNA synthesis were carried out by the same method described above. Primer sequences for *DGAT1* inside fragments and yeast *Actin1* as an internal reference gene are shown in Supplementary Table S2-2.

### Site-directed mutagenesis of GmDGAT1A

Single amino acid residue substitutions and site-saturation mutagenesis at sites 149, 205 or 263 were introduced into the native GmDGAT1A. Briefly, the full length of the native *GmDGAT1A* coding region was cloned into a pGEM^R^-T Easy vector (Promega, USA). Site-directed mutagenesis was performed by inverse PCR using PrimeSTAR GXL polymerase (Takara, Japan) and the pGEM^R^-T-vector as a template. The mutagenic primers were designed to change phenylalanine to leucine (F149L), phenylalanine to valine (F205V), or alanine to valine (A263V). These primer sequences are shown in Supplementary Table S2-2. The full length of the coding region in the pGEM^R^-T-vector was sequenced and single site mutagenesis was confirmed. The single site mutated *GmDGAT1A* variants were introduced into yeast H1246, and cultured as described above. The *HanDGAT1* introduced line was also cultured as an example of highly active DGAT1.

### Lipid Analysis

Lyophilized Arabidopsis mature seeds or five day induction-cultured yeast cells were lyophilized and triheptadecanoin (tri-17:0) was added at 10 µg/mg tissue as a standard. The samples were finely ground with a mortar and pestle with 2-3 mL of chloroform and methanol (2:1) containing 0.001% butylated hydroxytoluene (BHT). After brief centrifugation, the chloroform phase was transferred into a new glass tube. Samples were divided into two aliquots. One was used for thin layer chromatography (TLC) and the other directly for gas chromatography (GC) analysis.

TLC was applied to separate individual lipid classes. The chloroform extracts were loaded on lanes of LK6D Silica gel 60A TLC plates (GE healthcare Japan Co. Ltd, Tokyo, Japan), and the plates were put in a chamber with hexane: diethyl ether: acetic acid (80:20:1, v/v/v) containing 0.001% BHT. After finishing chromatography, the plate was dried and then sprayed with 0.005% primulin in 80% acetone, followed by visualizing under UV light. The TAG bands were scraped and transferred to a glass tube. The TAGs were eluted with 2 mL of diethyl ether three times.

For GC analysis, the samples were completely dried under an N_2_ stream and mixed with 0.5 mL of 18-mg/mL sodium methoxide (NaOCH_3_) in methanol then incubated for 45 min with shaking at room temperature. Each dried sample was resolved in 2 mL of hexane containing 0.001% BHT. Then, the hexane layer was extracted and concentrated under an N_2_ stream. Finally, the fatty acid methyl esters were analyzed with GC on a GC-14B (Shimazu Co. Ltd., Kyoto, Japan) with a 2.1 m × 0.25 mm glass column filled with Unisole 3000 (GL Sciences Co Ltd., Tokyo, Japan) and a flame ionization detector (GC-FID).

### Bioinformatics

Three dimension structures of VgDGAT1A, GmDGAT1A and GmDGAT1A-F146L were analyzed by I-TASSER (Yang and Zhang, 2015) and COFACTOR (Zhang *et al*., 2017) based on comparing with human DGAT1 (Wang *et al*., 2020).

## Results and discussion

### Extent of transgenic Arabidopsis seed oil increase were dependent on the DGAT1 origin

cDNAs of *AtDGAT1, GmDGAT1A*, castor *DGAT1* (*RcDGAT1*), and *VgDGAT1A* were introduced into *Arabidopsis thaliana* (Col-0) under the control of a seed specific promoter. Transgenic Arabidopsis lines were designated as AtD-OE (*AtDGAT1* over-expression), GmD (*GmDGAT1A*), RcD (*RcDGAT1*), and VgD (*VgDGAT1A*), respectively. Oil content and fatty acid composition of T3 seeds from transgenic lines were analyzed by GC. When two individual transgenic lines from each *DGAT1* introduced plant showed close oil contents, then their results were combined. All VgD lines showed a significantly higher oil content (average 44% relative increase compared to the wild type) followed by GmD (average 32% increase), RcD (average 22% increase), and AtD-OE lines showed little increase (Fig. 1A) even though the mRNA was expressed 30-fold higher than the wild-type (Supplementary Table S3). These results reflected our previous yeast micro assay study (Hatanaka *et al*., 2016). The fatty acid compositions did not change drastically between wild-type plants and transgenic lines (Fig. 1B). However, in GmD lines, there seems to be a tendency to use α-linolenic acid (18:3^Δ9,12,15^) more than oleic acid (18:1^Δ9^) as a substrate.

**Fig. 1.**
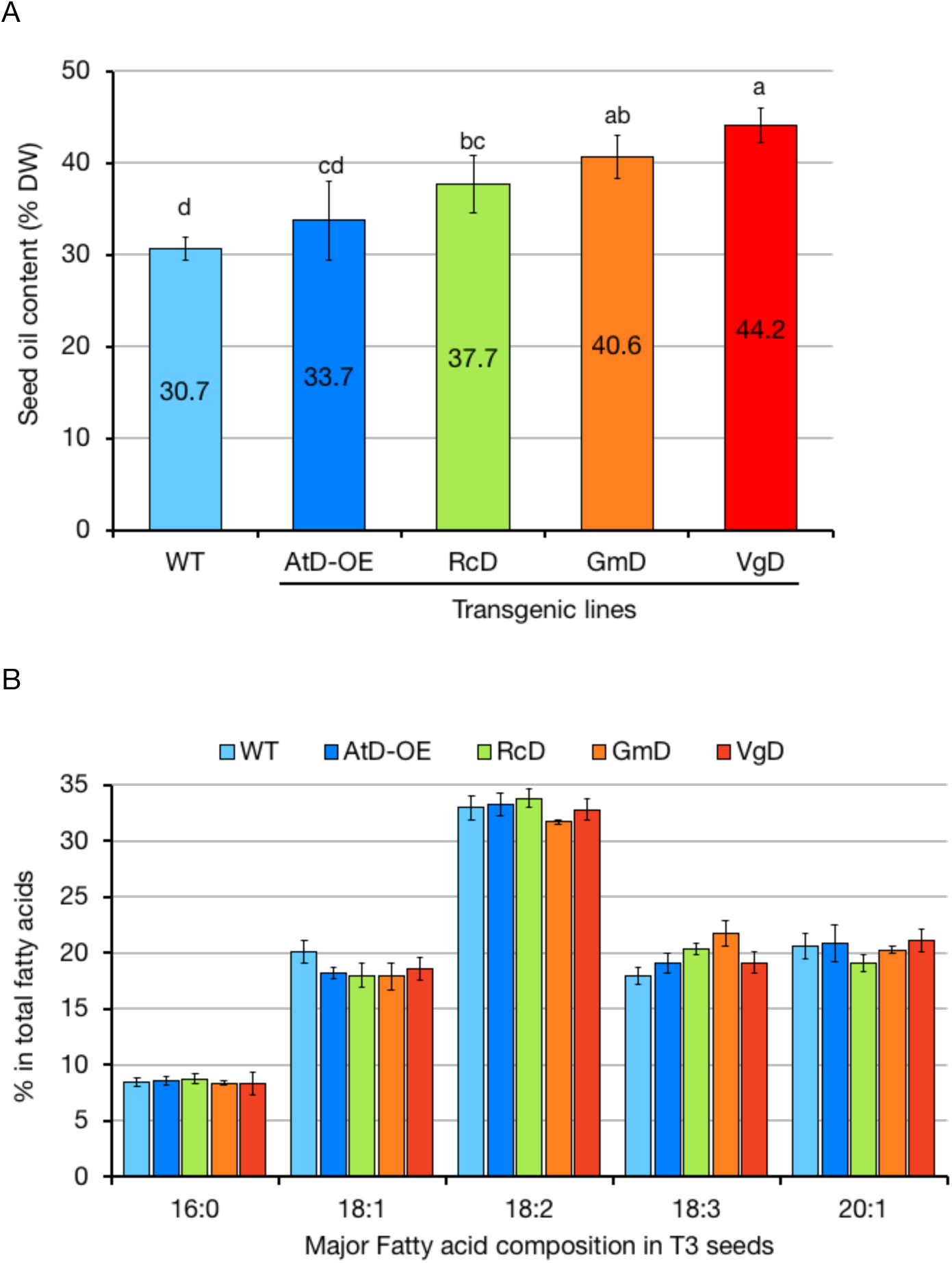
T3 seed oil contents (A) and the fatty acid compositions (B) of exogenous *DGAT1* gene expressing Arabidopsis lines. WT, wild-type; AtD-OE, *AtDGAT1* over-expression line; RcD, *RcDGAT1* expressing line; GmD, *GmDGAT1A* expressing line; VgD, *VgDGAT1A* expressing lines, respectively. Values indicate the mean ± SD of six plants. (A) T3 seed oil contents. *Different letters* indicate a significant difference (*p*<0.05) between transgenic lines based on Tukey’s HSD test. (B) Major fatty acid compositions of T3 seed oil. 16:0: palmitic acid; 18:1, oleic acid; 18:2, linoleic acid; 18:3, α-linolenic acid.

### In yeast H1246, DGAT1s from seven species were roughly classified into two groups

Addition to the four DGAT1s above, sunflower DGAT1 (*HanDGAT1*), Jatropha DGAT1 (*JcDGAT1*), and sesame DGAT1 (*SiDGAT1*) were introduced into the yeast quadruple knock-out strain H1246. Empty vector control transgenic yeast line were designated as ”VC”, “At” for *AtDGAT1*, “Rc” for *RcDGAT1*, “Gm” for *GmDGAT1A*, “Vg” for *VgDGAT1A*, “Ha” for *HanDGAT1*, “Jc” for *JcDGAT1*, and “Si” for *SiDGAT1*. Five day cultured cells were harvested and submitted to lipid analysis. The TLC results showed that they were obviously classified into a low TAG accumulating group (At, Rc, and Gm) and a high TAG accumulating group (Vg, Ha, Jc, and Si) (Supplementary Fig, S1A). We designated them “Low group” for the former three and “High group” for the latter four. Their total lipid and TAG contents were measured by GC (Fig. 2A). This result strongly support the TLC image in Fig. S1. All introduced gene expressions were confirmed by reverse-transcript PCR. The introduced genes were present even in the “Low group” lines (Supplementary Fig. S2). Their fatty acid compositions were also significantly different between two groups, e.g., the level of palmitic acid (16:0) was higher and that of oleic acid (18:1^Δ9^) was lower in the High group than the Low group (Fig. 2B). In yeast culture cells which accumulated a significant amount of TAG, the fatty acid compositions of total lipids reflected that of TAG. Differences in the amount of stearic acid (18:0) was a little higher in the High group than in the Low group. These differences suggested that the increased TAG in the High group has a different fatty acid composition compared to the total fatty acids of VC, which would consist mainly of polar lipids. Since no fatty acid or lipid was supplemented in the induction cultures, the distribution of fatty acids might be altered in the High group’s *DGAT1* introduced yeast cells.

**Fig. 2.**
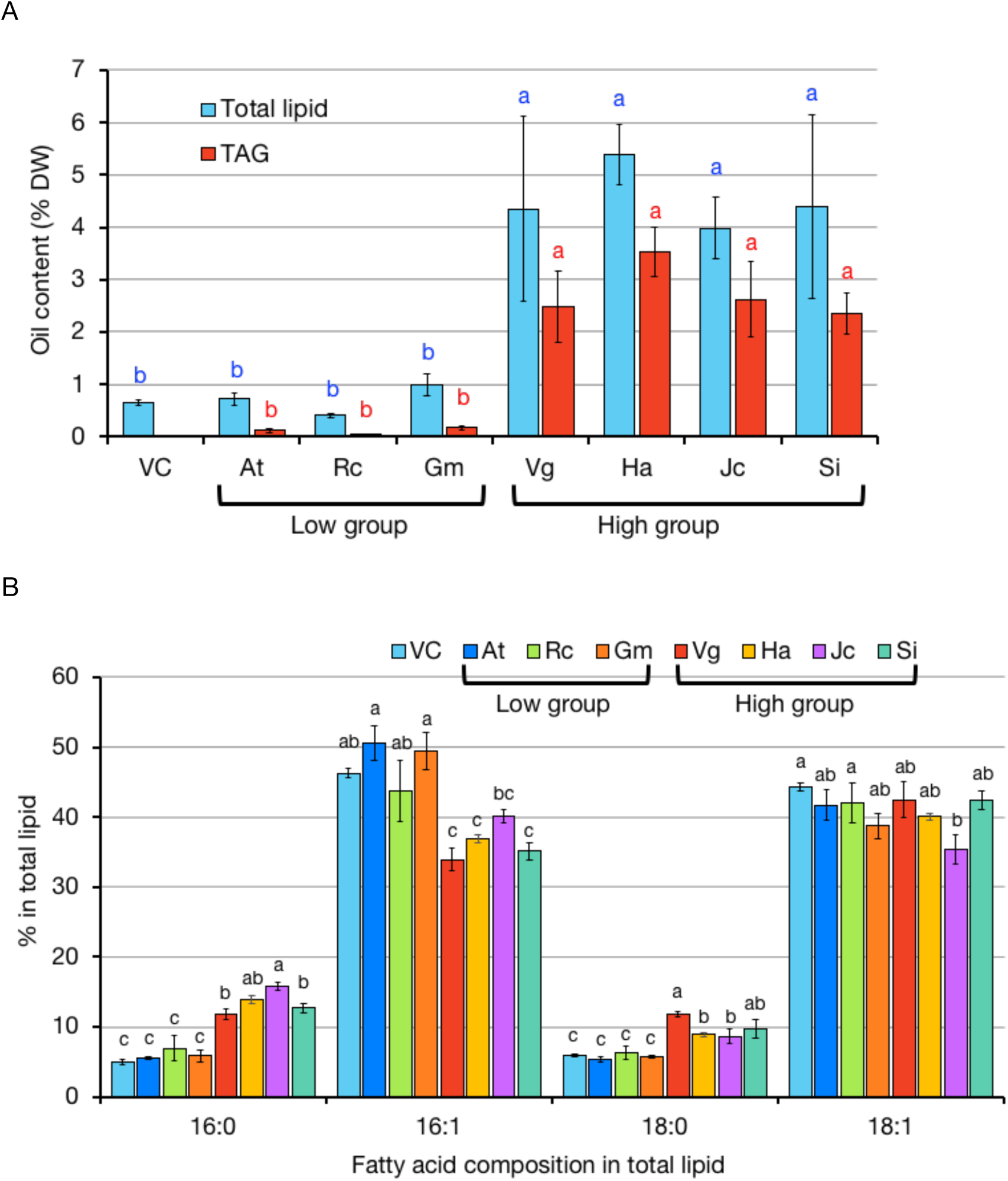
Oil content and fatty acid composition of yeast(H1246)transgenic lines measured by GC. (A) Total lipid (blue column) and TAG (red column) contents. (B) Fatty acid compositions in total lipid. 16:0, palmitic acid; 16:1, palmitoleic acid; 18:0, stearic acid; 18:1, oleic acid; 20:1, eicosenoic acid. VC, empty vector control; At, *AtDGAT1*; Rc, *RcDGAT1*; Gm, *GmDGAT1A*; Vg, *VgDGAT1A*; Ha, *HanDGAT1*; Jc, JcDGAT1; *Si, SiDGAT1* expressing lines, respectively. Values represent the means ± S.E. of 5 - 13 replications. There were significant differences between “Low group” and “High group” in both total lipid and in TAG based on Tukey’s HSD test (*p*<0.05).

### Three amino acid residues in the conserved region were different between the High and Low group

Fig. 3 shows the amino acid sequence alignment of the conserved region of the DGAT1s tested in the yeast expression culture and rapeseed BnaDGAT1. The sequence BnaDGAT1 is close to AtDGAT1 because they both belong to the Brassicaceae family. BnaDGAT1 is well studied (Chen *et al*., 2017), so it was added as a reference. The upper four DGAT1s in red letters were in the “High group”. A comparison between upper four and lower four, three amino acid sites were different in (1), (2), and (3) in Fig. 3. In VgDGAT1A of the “High group” leucine (L) was at site 174, valine (V) at 230, and valine (V) at 288, whereas they were phenylalanine (F) at site 149, phenylalanine (F) at 205, and alanine (A) 263 in GmDGAT1A of the “Low group”. Corresponding sites of the other proteins are listed in Supplementary Table S4. Jatropha and castor both belong to Euphorbiaceae family and their full length sequences are close (78.82% identity) whereas the three amino acid residues (1), (2), and (3) are different. Thus it may cause the difference of DGAT1 activity. These three amino acids have not been referred in previous reports to improve DGAT1s by amino acid substitution in GmDGAT1B (Roesler *et al*., 2016) and BnaDGAT1 (Chen *et al*., 2017).

**Fig. 3.**
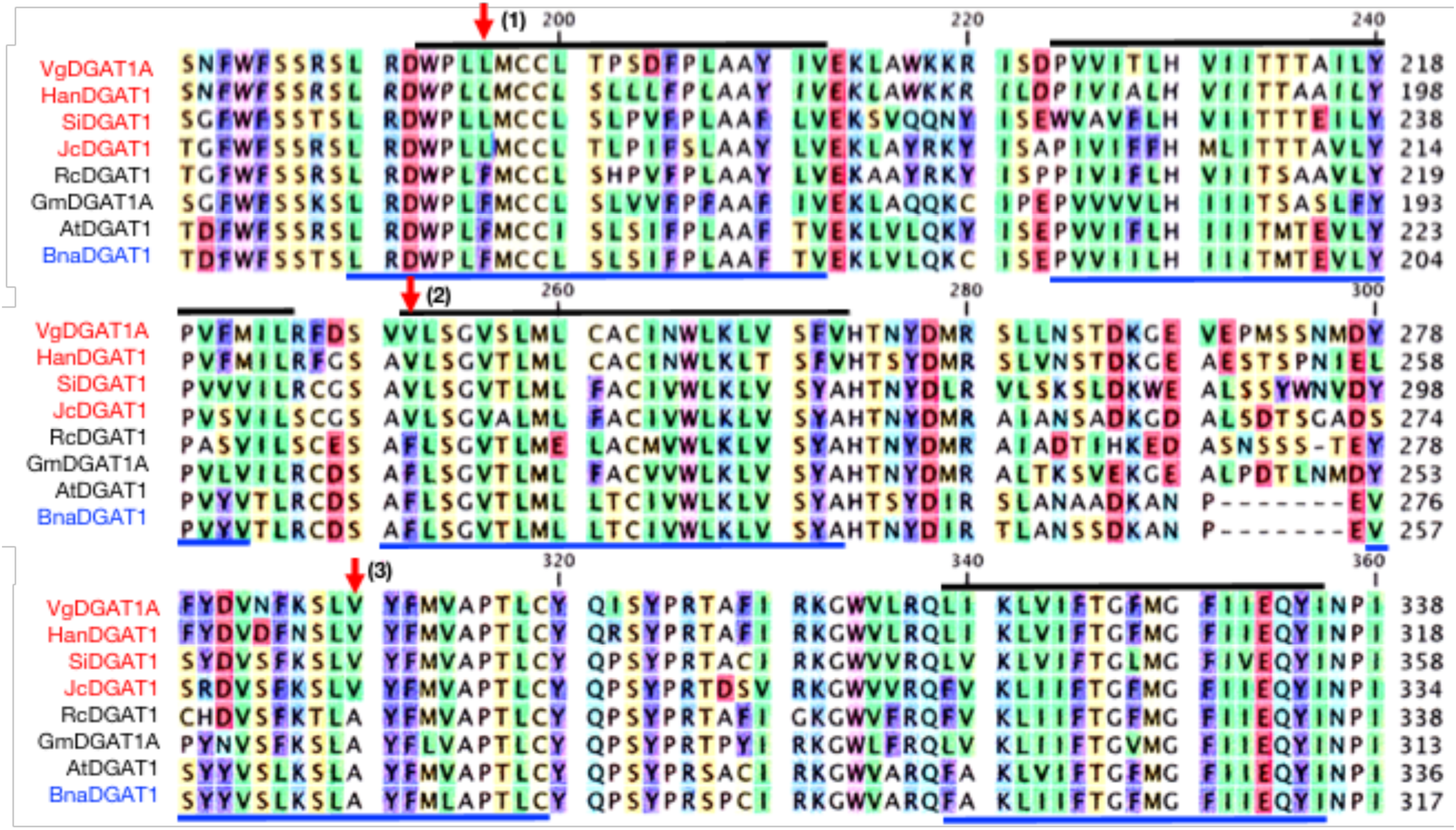
Amino acid sequence alignment of VgDGAT1A, HanDGAT1, SiDGAT1, JcDGAT1, RcDGAT1, AtDGAT1, GmDGAT1A and BnaDGAT1. Red arrows indicate the different amino acid residues between the upper four (High group) and the lower four. Black lines show trans-membrane domains (TMDs) of upper six DGAT1s and blue lines show TMDs of lower two DGAT1s of Brassicaceae family. TMDs were predicted by Phobius.

In other oil crop DGAT1s, tung tree (*Vercinia fordii*) DGAT1 (VfDGAT1) has the same three amino acid residues, oil palm (*Elaeis guineensis*) DGAT1 (EgDGAT1) has L at 171, I at 227, and V at 277, olive (*Olea europaea*) DGAT1 (OeDGAT1) has L at 183, V at 238, and A at 297, and linseed (*Linum usitatissimum*) DGAT1 (OeDGAT1) has F at 157, F at 213, and V at 271 (Supplementary Table S4). DGAT1s of olive and linseed may show medial characteristics between the “High” and the “Low” groups.

Black and blue lines in Fig. 3 indicate predicted trans-membrane helices (TMs) by Phobius. Most DGAT1s have nine TMs (black lines), however DGAT1s of the Brassicaceae family have 10 TMs (Blue lines). The amino acid residue (1) is in the second TM (TM2), (2) is in the fourth TM (TM4), and (3) is in the long loop between TM4 and TM5 (Fig. 4A), however, a region around (3) is within the membrane in Brassicaceae DGAT1s.

**Fig. 4.**
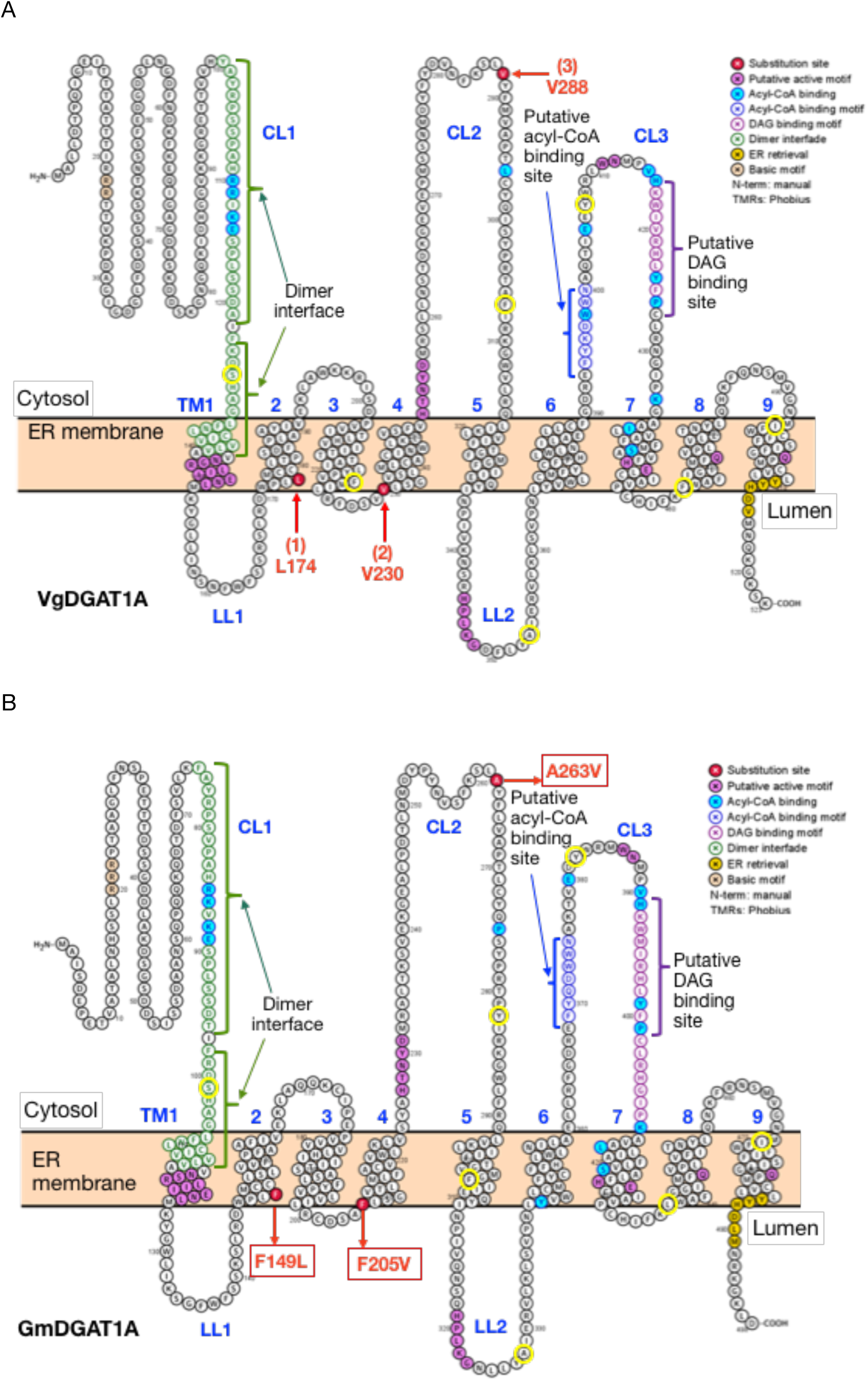
The predicted topology models of VgDGAT1A (A) and (B) GmDGAT1A created by Protter (Omasits *et al*., 2014). The trans-membrane domains were predicted by Phobius. Residues in color indicate that Red, residues for substitution; Pink, putative active site; Green, dimer forming interface; Blue, putative acyl-CoA binding site; Purple, putative DAG binding site. Yellow circles indicate amino acid residues tested in BnaDGAT1 (Chen *et al*., 2017). Putative functional sites are according to several reports (Chen *et al*., 2017; Sui *et al*., 2020; Wang *et al*., 2020). (A) VgDGAT1A. Red numbers and arrows are predicted key amino acid residues in this study. (B) GmDGAT1A. Site-directed amino acid substitutions are shown in red letters..

### A single amino acid substitution in GmDGAT1A increased TAG production

To further analyze the effect of amino acid residue substitutions of DGAT1 of Low group on storage lipid biosynthesis, site saturation mutagenesis was performed at site F149L, F205V and A263V in GmDGAT1A (Fig. 4B). All three GmDGAT1A variants introduced yeast H1246 lines accumulated higher TAG contents than the native GmDGAT1A introduced line, particularly F149L and A263V with 4.3 fold higher TAG in F149L 2,2 and 3.7 fold higher TAG in A263V (Fig. 5). Interestingly, substituted amino acid residues were all hydrophobic even the third site (A263V) that is predicted to locate on the loop between the fourth and TM5. In Brassicaceae DGAT1, this position would be within the TM domain. It may be possible that this site and adjacent amino acid residues are peripheral on or partially within the membrane. Chen *et al*. (2017) also reported that the replacement of I447 of BnaDGAT1 with A, C, F, L, T or V resulted in substantially higher neutral lipid content, whereas the replacement of I447 with D, E, N, R, K or Y resulted in lower neutral lipid content. As I, A, C, F, L and V are hydrophobic amino acid residues, polar amino acids such as D, E, N, R and K are inappropriate to exist within membranes. Almost all DGAT1s tested in this study have “I” at the corresponding site 447 of BnaDGAT1 except SiDGAT1 (M514) (Supplementary Table S5). Chen *et al*. (2017) also mentioned that L441P increased neutral lipid content. Among seven DGAT1s in this study, GmDGAT1A, AtDGAT1 RcDGAT1 (the Low group) and JcDGAT1 have “L”, whereas VgDGAT1A, HanDGAT1, SiDGAT1 (the High group) have “F” at site 441 of BnaDGAT1. L, I, F and P are all hydrophobic, however, “F” and “P” are aromatic amino acids that means they have a bulky benzene ring, but “L” and “I” do not (Fig. 4A). This may affect the peptide structure and F146L in GmDGAT1A in this study and I447P in BnaDGAT1 (Chen *et al*., 2017) increased TAG accumulation likely due to the structural alteration. In their study (2017), the majority of BnaDGAT1 variants with single amino acid residue substitutions that resulted in higher TAG accumulation, were either within or close to a TM domain. Exceptions were, C286Y on the loop between the TM5 and TM6, G332A on the loop between the sixth and the TM7, and Y386F on the loop between the TM7 and TM8. Looking at other DGAT1s, these three amino acids are commonly on the loop region. However, the corresponding amino acid residues of G332A of BnaDGAT1 was originally “A” in our tested seven DGAT1s. It may not be possible that hydrophobic amino acid residues work efficiently outside the membranes in DGAT1 activity.

**Fig. 5.**
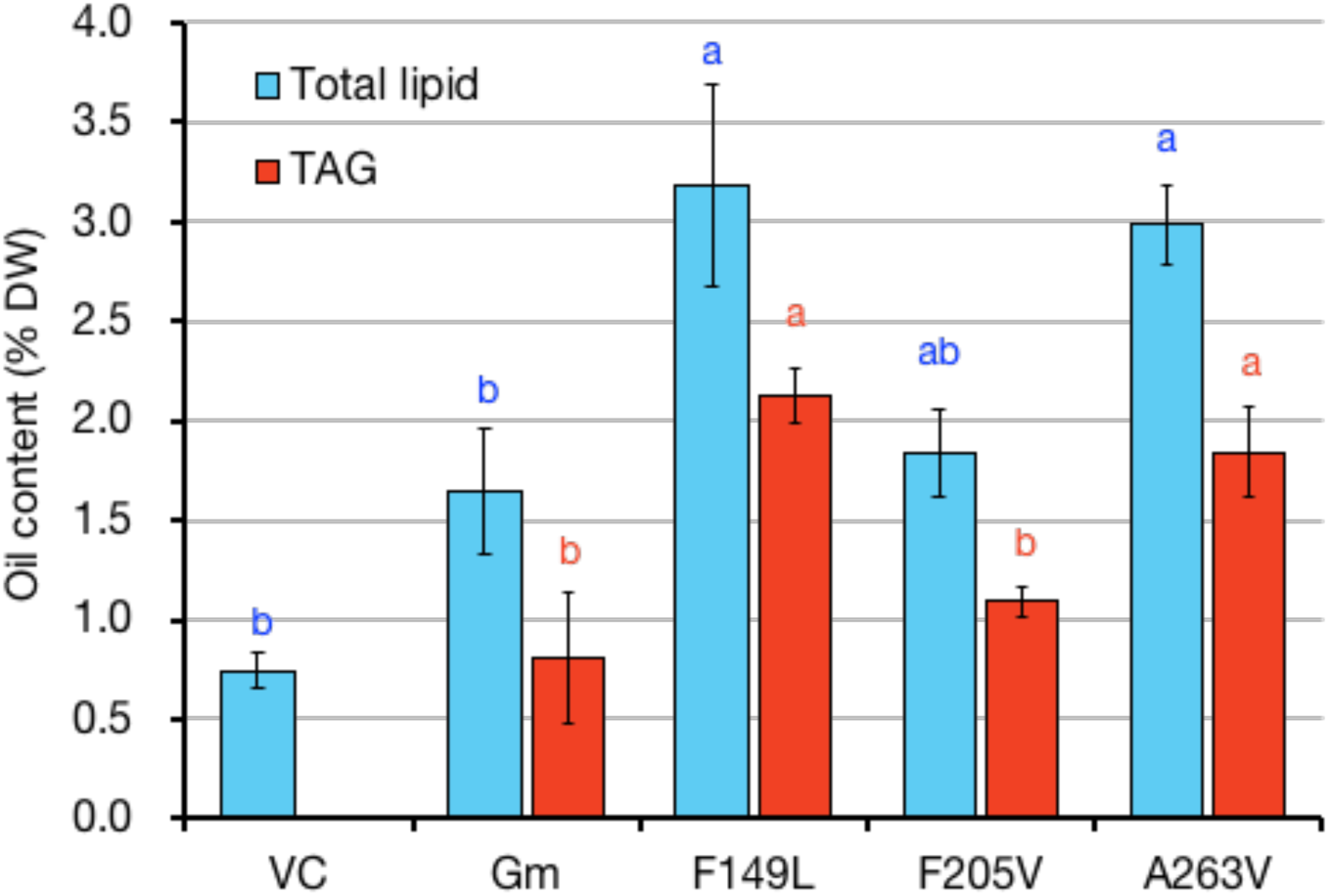
Total lipid (blue column) and TAG (red column) contents of yeast H1246 strains hosting *GmDGAT1A* variants generated by single-site mutagenesis. VC, empty vector control; Gm, native *GmDGAT1A*; F149L, F205V, A263V, *GmDGAT1A* variants. Oil contents of the yeast (H1246) cells were analyzed by GC-FID. *Different letters* indicate a significant difference (*p*<0.05) between variant lines based on Tukey’s HSD test (n = 5).

Surprisingly or interestingly, the double mutations (F149L/F205V, F205V/A263V, F149L/A263V) or the triple mutation (F149L/F205V/A263V) of GmDGAT1A were less effective in increasing TAG content than the single mutations (F149L or A263V) (data not shown). On the other hand, GmDGAT1b-MOD reported by Roesler *et al*. (2016) have 13 amino acid substitutions.

The fatty acid compositions of variants changed, particularly, the ratio of palmitoleic acid (16:1Δ9) significantly decreased and that of other fatty acids increased compared with the native GmDGAT1A (Fig 6A, B). This tendency was observed in DGAT1s of the High group (Fig. 2B). In general, DGAT1 does not have strong substrate specificities, however, it may have some preferences.

**Fig. 6.**
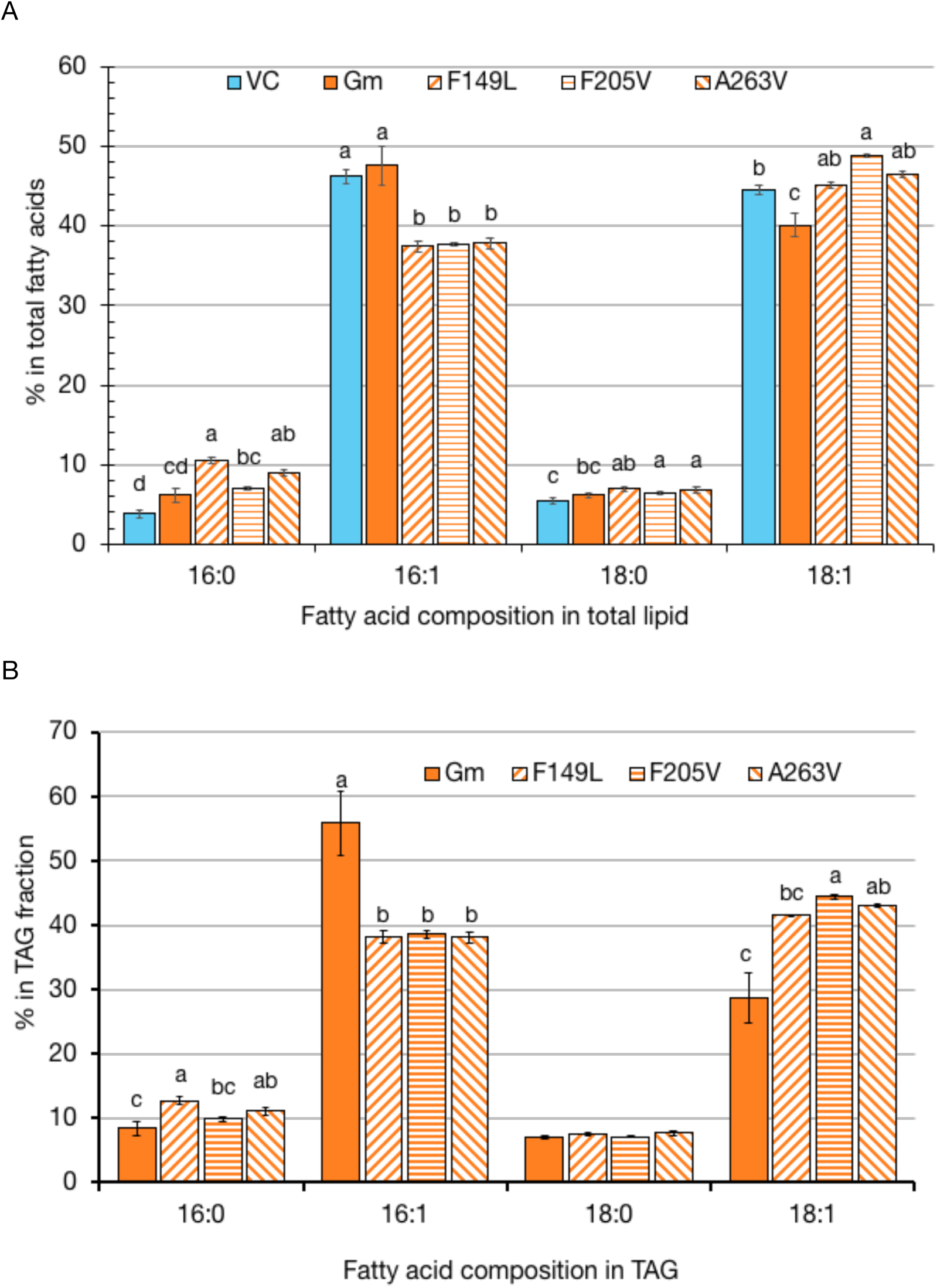
Fatty acid composition in total lipid (A) and TAG fraction (B) of yeast 679 H1246 strains hosting 680 *GmDGAT1A* variants generated by single-site mutagenesis. VC, empty vector control; Gm, native 681 *GmDGAT1A* (, *GmDGAT1A* variants (F149L, F205V and A263V). Fatty acids of the yeast cells were 682 analyzed by GC-FID. *Different letters* indicate a significant difference (*p*<0.05) between variant lines based 683 on Tukey’s HSD test (n = 5).

### Bioinformatics of DGAT1 proteins

I-TASSER (Yang and Zhang, 2015; Zhang *et al*., 2017) is a protein structure prediction and structure-based function annotation web-based server which was ranked as the No. 1 protein structure prediction server in the 14th CASP experiment (https://www.predictioncenter.org/casp14/zscores_final.cgi?gr_type=server_only) on December 2020. I-TASSER simulations generate decoys which are a large ensemble of structural conformations. Then I-TASSER uses the SPICKER program to cluster all the decoys based on the pair-wise structure similarity, and reports up to five models which corresponds to the five largest structure clusters. The confidence of each model is quantitatively measured by C-score (range of [-5, 2]) that is calculated based on the significance of threading template alignments and the convergence parameters of the structure assembly simulations. I-TASSER match the best confidence model with all structures in the Protein Data Bank (PDB) library by TM-align structural alignment program and scored them with TM-score. The best match model for VgDGAT1A, GmDGAT1A, and GmDGAT1A-F149L was human DGAT1 (Wang *et al*., 2020). TM-score for VgDGAT1A, GmDGAT1A, and GmDGAT1A-F149L were 0.762, 0.799, and 0.801, respectively. Due to these high scores, VgDGAT1A, GmDGAT1A, and GmDGAT1A-F149L proteins in this study have very similar structure to the human DGAT1 protein. Additionally, the web-based tool has predicted the function of the I-TASSER structure of the target protein through the COACH and COFACTOR programs. The COFACTOR deduces protein functions (ligand-binding sites, EC and GO) using structure comparison and protein-protein networks and COACH is a meta-server approach that combines multiple function annotation results (on ligand-binding sites) from the COFACTOR, TM-SITE and S-SITE programs. According to the prediction, the best ligand binding sites for VgDGAT1A was Manganese (2+) (with 1twaA PBD hit and confidence score: 0.09) at 277 and 281 ligand binding site residues (Fig. 6). The best ligand binding sites for GmDGAT1A was nucleic acid (with 4 ζ 6aA PBD hit and confidence score: 0.08) at 372, 375, 377, 380, and 382 ligand binding site residues (Fig. 7A). Also, The best ligand binding sites for GmDGAT1A-F149L was HEGA-10 (with 2y04B PBD hit and confidence score: 0.06) at 220 and 221 ligand binding site residues (Fig. 7B). HEGA-10 is a detergent used for solubilizing protein which has biomolecular interactions with voltage-gated sodium channel (McCusker *et al*., 2012). Interestingly, the predicted model for all were the same but the ligand-binding site were absolutely different (Figs. 7 and 8). This difference would be occurred by single site-directed mutagenesis of F149L for GmDGAT1A. These results are supporting our yeast experiment’s results that substituting this amino acid has a great role in changing protein function and might lead to produce more TAG.

**Fig. 7.**
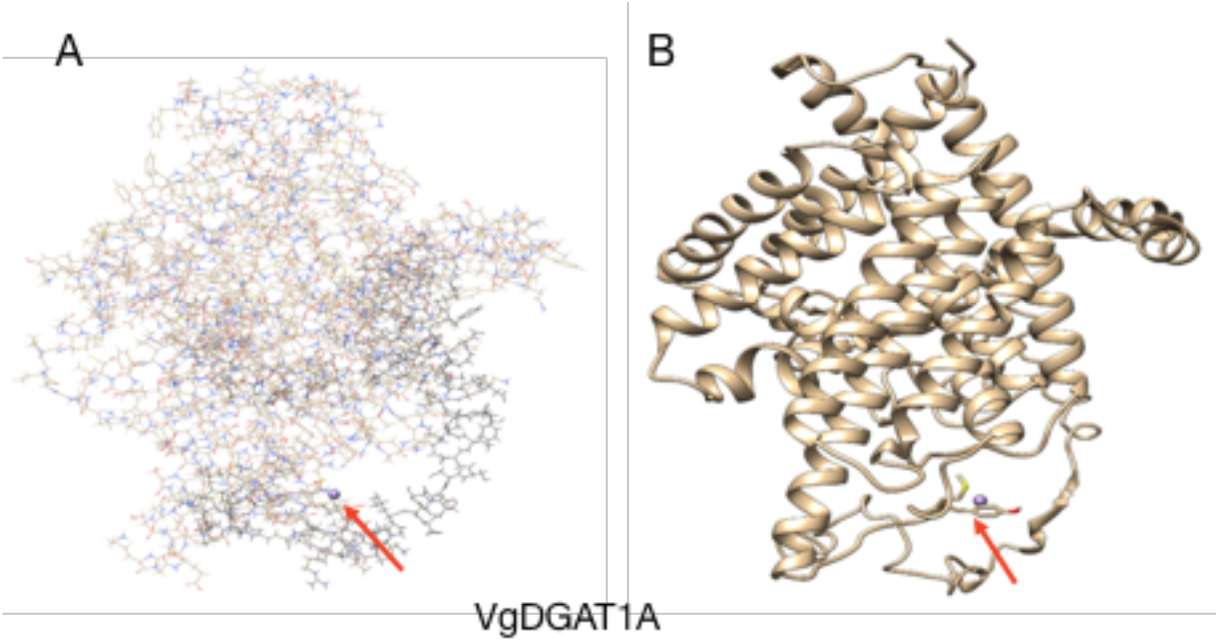
Two view pictures of ligand binding site residues for VgDGAT1A. Atomic view (A) and ribbon view (B) with manganese (2+) ligand binding site at 277 and 281 residues (red arrows).

**Fig. 8.**
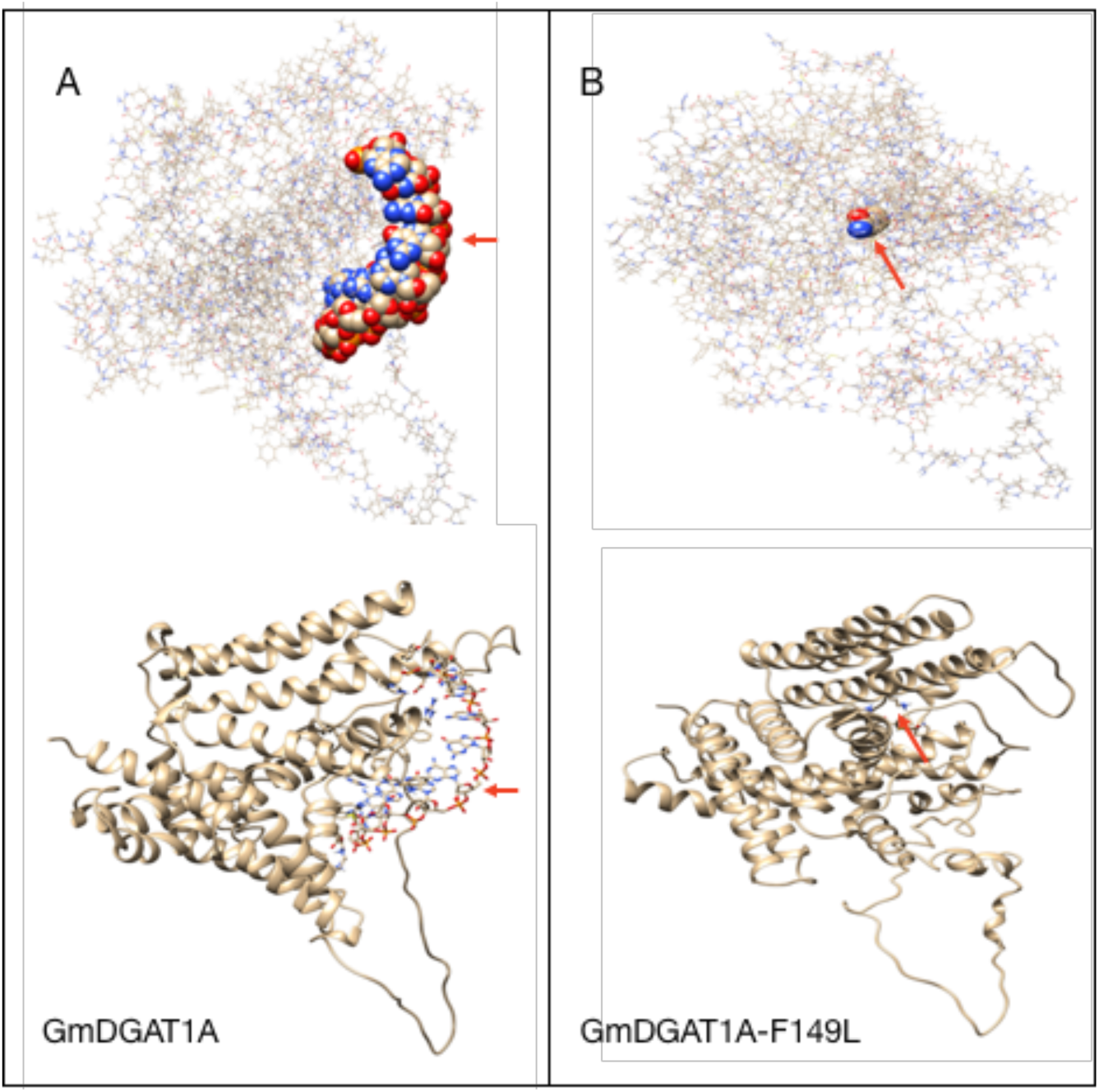
Two view pictures of ligand binding site residues for GmDGAT1A (A) and GmDGAT1A-F149L (B). A. Top, Atomic view; bottom, ribbon view of GmDGAT1A with nucleic acid ligand binding site at 372, 375, 377, 380 and 382 residues. B. Top, Atomic view; bottom, ribbon view of GmDGAT1A-F149L with HEGA-10 ligand binding site at 220 and 221 residues (red arrows).

DGAT1 belongs to the membrane-bound *O*-acyltransferase (MBOAT) superfamily, found in all kingdoms of life. DGAT1 from plants and mammals was previously shown to form a dimer (Caldo *et al*., 2017; McFie *et al*., 2010). This dimerization was confirmed by the recent cryo-electron microscopy (cryo-EM) studies of human DAGT1 (Sui *et al*., 2020; Wang *et al*., 2020). Each protomer of human DGAT1 has nine trans-membrane (TM) helices, TM1-TM9, with N and C termini located on the cytosolic and luminal sides of the endoplasmic reticulum membrane, respectively. Short helices in cytosolic loop (CL) and luminal loop (LL) regions orient in parallel to the membrane surface (Wang 2020). This structure is very similar to topology models of VgDGAT1A and GmDGAT1A (Fig. 4). Crossover of the TM1 helix allows the N-terminal CL1 of one protomer to interact with both CL1 and CL2 of the another protomer (Sui *et al*., 2020). TM2-TM9 and the two cytosolic loops, CL2 and CL3 form a distinctive structural fold that called the MBOAT fold (Wang *et al*., 2020) (Fig. 9). The MBOAT fold of DGAT1 carves out a large hollow chamber within the membrane that is open to the bilayer via a wide lateral gate and this region is considered as DGAT1 active sites (Sui *et al*., 2020; Wang *et al*., 2020). The three amino acid residues in this study face to this reaction chamber (Fig. 9).

**Fig. 9.**
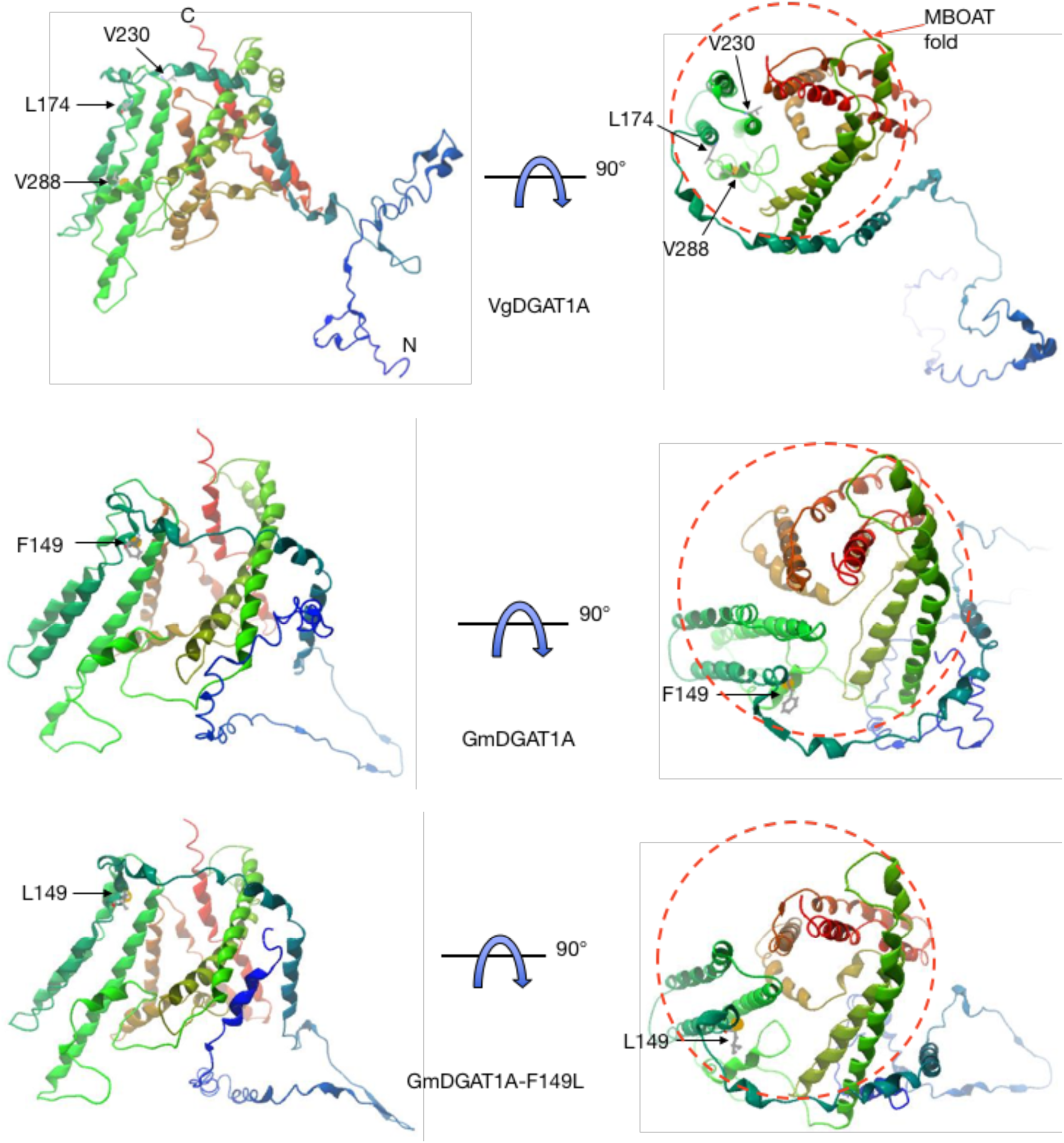
Three-dimensional structures of VgDGAT1A, GmDGAT1A, and GmDGAT1A-F149L. Cartoons represent each DGAT1 protomer in two orientations. The MBOAT fold is marked with a red dashed circle.

In conclusion, our results show that three amino acids, L174, V230 and V288 in VgDGAT1A, 154L, 210V and 268V in HanDGAT1, 194L, 250V and 308V in SiDGAT1, 154L, 226V and 284V in JcDGAT1, can be key factors of the High group’s DGAT1 activity (Supplementary Table S4). Also, considering examples of BnaDGAT1 variants (Chen *et al*., 2017), these results support that only single amino acid substitution are effective in increasing activities of low performance DGAT1s. As we show the primer sequences for site-directed mutagenesis (Supplementary Table S2), only one base substitution is needed to provide an improved DGAT1. This technique may be applied to a genome-editing method easily.

Amino acid substitution experiments of AtDGAT1 are ongoing and the results to date are even more promising than GmDGAT1A variants.

## Supplementary data

The following supplementary data are available at *JXB* online.

*Table S1*. GenBank numbers of genes mentioned in this study.

*Table S2*. Primer sequences used in the current study.

*Table S3*. Quantitative gene expression analysis in transgenic Arabidopsis siliques.

*Table S4*. Corresponding amino acid residue sites that were numbered on Fig. 3 in various plant DGAT1.

*Table S5*. Corresponding amino acid residue sites that were tested in BnaDGAT1 (data from Chen *et al*. 2017).

*Fig. S1*. An example of TLC development of total lipid extract from transgenic yeast lines (H1246).

*Fig. S2*. Reverse-transcript-PCR of transgenic yeast lines (H1246).

*Fig. S3*. An example of TLC development of total lipid extract from transgenic yeast lines (H1246) expressing *GmDGAT1A* variants.

## Acknowledgement

The authors would like to thank Prof. Sten Stymnes (Department of Plant Breeding, Swedish University of Agricultural Sciences, Alnarp, Sweden) for providing *S. cerevisiae* H1246 strain.

## Author contributions

TH designed and performed most of the experiments, analyzed the data and prepared the initial draft of the manuscript; YT performed Arabidopsis transformation and obtained all transgenic lines in this study; DM supervised yeast experiments; HF helped vector constructions and other molecular techniques, DS and TA contributed to valuable discussions during study; MS worked on bioinformatics; DH supervised this whole study and manuscript editing. All authors contributed to the preparation of the final article.

## Data availability

All relevant data can be found within the manuscript and its supplementary data.

## References

Anderson MA, Collier L, Dilliplane R, Ayorinde FO. 1993. Mass Spectrometric characterization of Vernonia galamensis oil. Journal of American Oil Chemists’ Society 70, 905–908.

Andrianov V, Borisjuk N, Pogrebnyak N, Brinker A, Dixon J, Spitsin S, Flynn J, Matyszczuk P, Andryszak K, Laurelli M, Golovkin M, Koprowski H. 2010. Tobacco as a production platform for biofuel: overexpression of Arabidopsis DGAT and LEC2 genes increases accumulation and shifts the composition of lipids in green biomass. Plant Biotechnology Journal 8, 277–287.

Ayorinde FO, Osman G, Shepard RL, Powers FT. 1988. Synthesis of azelaic acid and suberic acid from Vernonia galamensis oil. Journal of American Oil Chemists’ Society 65, 1774–1777.

Banilas G, Karampelias M, Makariti I, Kourti A, Hatzopoulos P. 2011. The olive DGAT2 gene is developmentally regulated and shares overlapping but distinct expression patterns with DGAT1. Journal of Experimental Botany 62, 521–532.

Beisson F, Koo AJ, Ruuska S, Schwender J, Pollard M, Thelen JJ, Paddock T, Salas JJ, Savage L, Milcamps A, Mhaske VB, Cho Y, Ohlrogge JB. 2003. Arabidopsis genes involved in acyl lipid metabolism. A 2003 census of the candidates, a study of the distribution of expressed sequence tags in organs, and a web-based database. Plant Physiology 132, 681–697.

Bouvier-Nave P, Benveniste P, Oelkers P, Sturley SL, Hubert S. 2000. Expression in yeast and tobacco of plant cDNAs encoding acyl CoA:diacylglycerol acyltransferase. European Journal of Biochemstry 267, 85–96.

Burgal J, Shockey J, Lu C, Dyer J, Larson T, Graham I, Browse J. 2008. Metabolic engineering of hydroxy fatty acid production in plants: RcDGAT2 drives dramatic increases in ricinoleate levels in seed oil. Plant Biotechnology Journal 6, 819–831.

Bustos MM, Iyer M, Gagliardi SJ. 1998. Induction of a ß-phaseolin promoter by exogenous abscisic acid in tobacco: developmental regulation and modulation by external sucrose and Ca^2+^ ions. Plant Molecular Biology 37, 265–274.

Caldo KMP, Acedo JZ, Panigrahi R, Vederas JC, Weselake RJ, Lemieux MJ. 2017. Diacylglycerol acyltransferase 1 is regulated by Its N-terminal domain in response to allosteric effectors. Plant Physiology 175, 667–680.

Carlson KD, Chang SP. 1985. Chemical eposidation of natural unsaturated epoxy seed oil from Vernonia galamensis and a look at epoxy oil markets. Journal of American Oil Chemists’ Society 62, 934–939.

Carlsson AS. 2009. Plant oils as feedstock alternatives to petroleum - A short survey of potential oil crop platforms. Biochimie 91, 665–670.

Carlsson AS, Yilmaz JL, Green AG, Stymne S, Hofvander P. 2011. Replacing fossil oil with fresh oil - with what and for what? European Journal of Lipid Science and Technology 113, 812–831.

Cases S, Smith SJ, Zheng YW, Myers HM, Lear SR, Sande E, Novak S, Collins C, Welch CB, Lusis AJ, Erickson SK, Farese RV. 1998. Identification of a gene encoding an acyl CoA: diacylglycerol acyltransferase, a key enzyme in triacylglycerol synthesis. Proceedings of the National Academy of Sciences, USA 95, 13018–13023.

Cases S, Stone SJ, Zhou P, Yen E, Tow B, Lardizabal KD, Voelker T, Farese RV, Jr. 2001. Cloning of DGAT2, a second mammalian diacylglycerol acyltransferase, and related family members. Journal of Biological Chemistry 276, 38870–38876.

Chen G, Xu Y, Siloto RMP, Caldo KMP, Vanhercke T, Tahchy AE, Niesner N, Chen Y, Mietkiewska E, Weselake RJ. 2017. High-performance variants of plant diacylglycerol acyltransferase 1 generated by directed evolution provide insights into structure function. Plant Journal 92, 167–177.

Clough SJ, Bent AF. 1998. Floral dip: a simplified method for Agrobacterium-mediated transformation of Arabidopsis thaliana. Plant Journal 16, 735–743.

Dahlqvist A, Ståhl U, Lenman M, Banas A, Lee M, Sandager L, Ronne H, Stymne S. 2000. Phospholipid:diacylglycerol acyltransferase: An enzyme that catalyzes the acyl-CoA-independent formation of triacylglycerol in yeast and plants. Proceedings of the National Academy of Sciences, USA 97, 6487–6492.

Fraser T, Waters A, Chatrattanakunchai S, Stobart K. 2000. Does triacylglycerol biosynthesis require diacylglycerol acyltransferase (DAGAT)? Biochemical Society Transactions 28, 698–700.

Gamborg OL, Miller RA, Ojima K. 1968. Nutrient requirements of suspension cultures of soybean root cells. Experimental Cell Research 50, 151–158.

Hatanaka T, Serson W, Li RZ, Armstrong P, Yu KS, Pfeiffer T, Li XL, Hildebrand D. 2016. A Vernonia diacylglycerol acyltransferase can increase renewable oil production. Journal of Agricultural and Food Chemistry 64, 7188–7194.

Hatanaka T, Yu K, Hildebrand DF. 2003. Cloning and expression of a Vernonia and Euphorbia diacylglycerol acyltransferase cDNAs. In: Murata N, Yamada M, Nishida I, Okuyama H, Sekiya J, w H, eds. Advanced research on plant lipids. Dordrecht, The Netherlands: Kluwer Academic Publishers, 155–158.

He XH, Turner C, Chen GQ, Lin JT, McKeon TA. 2004. Cloning and characterization of a cDNA encoding diacylglycerol acyltransferase from castor bean. Lipids 39, 311–318.

Hernandez ML, Whitehead L, He Z, Gazda V, Gilday A, Kozhevnikova E, Vaistij FE, Larson TR, Graham IA. 2012. A cytosolic acyltransferase contributes to triacylglycerol synthesis in sucrose-rescued Arabidopsis seed oil catabolism mutants. Plant Physiology 160, 215–225.

Hobbs DH, Lu CF, Hills MJ. 1999. Cloning of a cDNA encoding diacylglycerol acyltransferase from Arabidopsis thaliana and its functional expression. FEBS Letters 452, 145–149.

Jako C, Kumar A, Wei Y, Zou J, Barton DL, Giblin EM, Covello PS, Taylor DC. 2001. Seed-specific over-expression of an Arabidopsis cDNA encoding a diacylglycerol acyltransferase enhances seed oil content and seed weight. Plant Physiology 126, 861–874.

Jaworski J, Cahoon EB. 2003. Industrial oils from transgenic plants. Current Opinion in Plant Biology 6, 178–184.

Kalscheuer R, Steinbuchel A. 2003. A novel bifunctional wax ester synthase/acyl-CoA:diacylglycerol acyltransferase mediates wax ester and triacylglycerol biosynthesis in Acinetobacter calcoaceticus ADP1. Journal of Biological Chemistry 278, 8075–8082.

Kawagoe Y, Campell BR, Murai N. 1994. Synergism between CACGTG (G-box) and CSCCTG cis-elements is required for activation of the bean seed storage protein ß-phaseolin gene. Plant Journal 5, 885–890.

King A, Nam JW, Han J, Hilliard J, Jaworski JG. 2007. Cuticular wax biosynthesis in petunia petals: cloning and characterization of an alcohol-acyltransferase that synthesizes wax-esters. Planta 226, 381–394.

Kroon JT, Wei W, Simon WJ, Slabas AR. 2006. Identification and functional expression of a type 2 acyl-CoA:diacylglycerol acyltransferase (DGAT2) in developing castor bean seeds which has high homology to the major triglyceride biosynthetic enzyme of fungi and animals. Phytochemistry 67, 2541–2549.

Lardizabal K, Effertz R, Levering C, Mai J, Pedroso MC, Jury T, Aasen E, Gruys K, Bennett K. 2008. Expression of Umbelopsis ramanniana DGAT2A in seed increases oil in soybean. Plant Physiology 148, 89–96.

Lardizabal KD, Mai JT, Wagner NW, Wyrick A, Voelker T, Hawkins DJ. 2001. DGAT2 is a new diacylglycerol acyltransferase gene family: purification, cloning, and expression in insect cells of two polypeptides from Mortierella ramanniana with diacylglycerol acyltransferase activity. Journal of Biological Chemistry 276, 38862–38869.

Li F, Wu X, Lam P, Bird D, Zheng H, Samuels L, Jetter R, Kunst L. 2008. Identification of the wax ester synthase/acyl-coenzyme A: diacylglycerol acyltransferase WSD1 required for stem wax ester biosynthesis in Arabidopsis. Plant Physiology 148, 97–107.

Li R, Hatanaka T, Yu K, Wu Y, Fukushige H, Hildebrand D. 2013. Soybean oil biosynthesis: role of diacylglycerol acyltransferases. Functional and Integrative Genomics 13, 99–113.

Li R, Yu K, Hatanaka T, Hildebrand DF. 2010a. Vernonia DGATs increase accumulation of epoxy fatty acids in oil. Plant Biotechnology Journal 8, 184–195.

Li R, Yu K, Hildebrand DF. 2010b. DGAT1, DGAT2 and PDAT expression in seeds and other tissues of epoxy and hydroxy fatty acid accumulating plants. Lipids 45, 145–157.

Li R, Yu K, Wu Y, Tateno M, Hatanaka T, Hildebrand DF. 2012. Vernonia DGATs can complement the disrupted oil and protein metabolism in epoxygenase-expressing soybean seeds. Metabolic Engineering 14, 29–38.

Lung S-C, Weselake RJ. 2006. Diacylglycerol acyltransferase: A key mediator of plant triacylglycerol Ssynthesis. Lipids 41, 1073–1088.

McCusker EC, Bagneris C, Naylor CE, Cole AR, D’Avanzo N, Nichols CG, Wallace BA. 2012. Structure of a bacterial voltage-gated sodium channel pore reveals mechanisms of opening and closing. Nature Communications 3, 1102.

McFie PJ, Stone SL, Banman SL, Stone SJ. 2010. Topological orientation of acyl-CoA:diacylglycerol acyltransferase-1 (DGAT1) and identification of a putative active site histidine and the role of the n terminus in dimer/tetramer formation. Journal of Biological Chemistry 285, 37377–37387.

Misra A, Khan K, Niranjan A, Nath P, Sane VA. 2013. Over-expression of JcDGAT1 from Jatropha curcas increases seed oil levels and alters oil quality in transgenic Arabidopsis thaliana. Phytochemistry 96, 37–45.

Murashige T, Skoog F. 1962. A revised medium for rapid growth and bio assays with tobacco tissue cultures. Physiologia Plantarum 15, 473–497.

Neff WE, Adlof RO, Konishi H, Weisleder D. 1993. High-performance liquid chromatography of the triacylglycerols of Vernonia galamensis and Crepis alpina seed oils. Journal of American Oil Chemists’ Society 70, 449–455.

Nykiforuk CL, Tara L. Furukawa-Sto¡er, Phillip W. Hu¡, Magdalena Sarna, Andrë Laroche, Maurice M. Moloney, Weselake RJ. 2002. Characterization of cDNAs encoding diacylglycerol acyltransferase from cultures of Brassica napus and sucrose-mediated induction of enzyme biosynthesis. Biochimica et Biophysica Acta 1580, 95–109.

Oelkers P, Tinkelenberg A, Erdeniz N, Cromleyi D, Billheimeri JT, Sturley SL. 2000. A lecithin cholesterol acyltransferase-like gene mediates diacylglycerol esterification in yeast. Journal of Biological Chemistry 275, 15609–15612.

Ohlrogge J, Browse J. 1995. Lipid biosynthesis. Plant Cell 7, 957–970.

Ohlrogge JB. 1994. Design of new plant products: Engineering of fatty acid metabolism. Plant Physiology 104, 821–826.

Omasits U, Ahrens CH, Muller S, Wollscheid B. 2014. Protter: interactive protein feature visualization and integration with experimental proteomic data. Bioinformatics 30, 884–886.

Perdue REJ, Carlson KD, Gilbert MG. 1986. Vernonia galamensis, potential new crop source of epoxy acid. Economic Botany 40, 54–68.

Rao SS, Hildebrand D. 2009. Changes in oil content of transgenic soybeans expressing the yeast SLC1 gene. Lipids 44, 945–951.

Roesler K, Shen B, Bermudez E, Li C, Hunt J, Damude HG, Ripp KG, Everard JD, Booth JR, Castaneda L, Feng L, Meyer K. 2016. An improved variant of soybean type 1 diacylglycerol acyltransferase increases the oil content and decreases the soluble carbohydrate content of soybeans. Plant Physiology 171, 878–893.

Rosli R, Chan PL, Chan KL, Amiruddin N, Low EL, Singh R, Harwood JL, Murphy DJ. 2018. In silico characterization and expression profiling of the diacylglycerol acyltransferase gene family (DGAT1, DGAT2, DGAT3 and WS/DGAT) from oil palm, Elaeis guineensis. Plant Science 275, 84–96.

Saha S, Enugutti B, Rajakumari S, Rajasekharan R. 2006. Cytosolic triacylglycerol biosynthetic pathway in oilseeds. Molecular cloning and expression of peanut cytosolic diacylglycerol acyltransferase. Plant Physiology 141, 1533–1543.

Sandager L, Gustavsson MH, Stahl U, Dahlqvist A, Wiberg E, Banas A, Lenman M, Ronne H, Stymne S. 2002. Storage lipid synthesis is non-essential in yeast. Journal of Biological Chemistry 277, 6478–6482.

Sanjaya, Miller R, Durrett TP, Kosma DK, Lydic TA, Muthan B, Koo AJ, Bukhman YV, Reid GE, Howe GA, Ohlrogge J, Benning C. 2013. Altered lipid composition and enhanced nutritional value of Arabidopsis leaves following introduction of an algal diacylglycerol acyltransferase 2. Plant Cell 25, 677–693.

Shockey JM, Gidda SK, Chapital DC, Kuan JC, Dhanoa PK, Bland JM, Rothstein SJ, Mullen RT, Dyer JM. 2006. Tung tree DGAT1 and DGAT2 have nonredundant functions in triacylglycerol biosynthesis and are localized to different subdomains of the endoplasmic reticulum. Plant Cell 18, 2294–2313.

Siloto RM, Truksa M, Brownfield D, Good AG, Weselake RJ. 2009. Directed evolution of acyl-CoA:diacylglycerol acyltransferase: development and characterization of Brassica napus DGAT1 mutagenized libraries. Plant Physiology and Biochemistry 47, 456–461.

Stahl U, Carlsson AS, Lenman M, Dahlqvist A, Huang B, Banas W, Banas A, Stymne S. 2004. Cloning and functional characterization of a phospholipid:diacylglycerol acyltransferase from Arabidopsis. Plant Physiology 135, 1324–1335.

Stobart K, Manuel Mancha, Marit Lenman, Anders Dahlqvist, Stymne S. 1997. Triacylglycerols are synthesised and utilized by transacylation reactions in microsomal preparations of developing sa, fflower (Carthamus tinctorius L.) seeds. Planta 203, 58–66.

Sui X, Wang K, Gluchowski NL, Elliott SD, Liao M, Walther TC, Farese RV, Jr. 2020. Structure and catalytic mechanism of a human triacylglycerol-synthesis enzyme. Nature 581, 323–328.

Taylor DC, Zhang Y, Kumar A, Francis T, Giblin EM, Barton DL, Ferrie JR, Laroche A, Shah S, Zhu W, Snyder CL, Hall L, Rakow G, Harwood JL, Weselake RJ. 2009. Molecular modification of triacylglycerol accumulation by over-expression of DGAT1 to produce canola with increased seed oil content under field conditions. Botany 87, 533–543.

Wang L, Qian H, Nian Y, Han Y, Ren Z, Zhang H, Hu L, Prasad BVV, Laganowsky A, Yan N, Zhou M. 2020. Structure and mechanism of human diacylglycerol O-acyltransferase 1. Nature 581, 329–332.

Wang Z, Huang W, Chang J, Sebastian A, Li Y, Li H, Wu X, Zhang B, Meng F, Li W. 2014. Overexpression of SiDGAT1, a gene encoding acyl-CoA:diacylglycerol acyltransferase from Sesamum indicum L. increases oil content in transgenic Arabidopsis and soybean. Plant Cell, Tissue and Organ Culture 119, 399–410.

Xu JY, Kazachkov M, Jia YH, Zheng ZF, Zou JT. 2013. Expression of a type 2 diacylglycerol acyltransferase from Thalassiosira pseudonana in yeast leads to incorporation of docosahexaenoic acid beta-oxidation intermediates into triacylglycerol. FEBS Journal 280, 6162–6172.

Xu Y, Pan X, Lu J, Wang J, Shan Q, Stout J, Chen G. 2021. Evolutionary and biochemical characterization of a Chromochloris zofingiensis MBOAT with wax synthase and diacylglycerol acyltransferase activity. Journal of Experimental Botany 72, 5584–5598.

Yang J, Zhang Y. 2015. I-TASSER server: new development for protein structure and function predictions. Nucleic Acids Research 43, W174–181.

Yu K, Li R, Hatanaka T, Hildebrand D. 2008. Cloning and functional analysis of two type 1 diacylglycerol acyltransferases from Vernonia galamensis. Phytochemistry 69, 1119–1127.

Yu KS, McCracken CT, Li RZ, Hildebrand DF. 2006. Diacylglycerol acyltransferases from Vernonia and Stokesia prefer substrates with vernolic acid. Lipids 41, 557–566.

Zhang C, Freddolino PL, Zhang Y. 2017. COFACTOR: improved protein function prediction by combining structure, sequence and protein-protein interaction information. Nucleic Acids Research 45, W291–W299.

Zou J, Wei Y, Jaco C, Kumar A, Selvaraj G, Taylor DC. 1999. The Arabidopsis thaliana TAG1 mutant has a mutation in a diacylglycerol acyltransferase gene. Plant Journal 19, 645–653.

